# Benefits of individualized brain anatomies and EEG electrode positions for auditory cortex localization

**DOI:** 10.1101/2022.06.15.496307

**Authors:** Karolina Ignatiadis, Roberto Barumerli, Brigitta Tóth, Robert Baumgartner

## Abstract

Due to its high temporal resolution and non-invasive nature, electroencephalography (EEG) is considered a method of great value for the field of auditory cognitive neuroscience. In performing source space analyses, localization accuracy poses a bottleneck, which precise forward models based on individualized attributes such as subject anatomy or electrode locations aim to overcome. Yet acquiring anatomical images or localizing EEG electrodes requires significant additional funds and processing time, making it an oftentimes inaccessible asset. Neuroscientific software offers template solutions, on which analyses can be based. For localizing the source of auditory evoked responses, we here compared the results of employing such template anatomies and electrode positions versus the subject-specific ones, as well as combinations of the two. All considered cases represented approaches commonly used in electrophysiological studies. We considered differences between two commonly used inverse solutions (dSPM, sLORETA) and targeted the primary auditory cortex; a notoriously small cortical region that is located within the lateral sulcus, thus more prone to errors in localization. Through systematical comparison of the outcomes in terms of auditory activity attributes and leakage quantification, we assessed how the individualization steps impacted the analyses outcomes. Both electrode locations as well as subject anatomies were found to have an effect, which though varied based on the configuration considered. When comparing the inverse solutions, we moreover found that dSPM more consistently benefited from individualization of subject characteristics. Based on the scientific question considered, our results may be used to facilitate the planning of auditory neuroscientific experiments in terms of expected infrastructure, personnel and funds.

## 1 INTRODUCTION

Being inexpensive and non-invasive, electroencephalography (EEG) is a widely used neuroimaging method. Due to its high temporal resolution it can reflect fast processes, which is especially advantageous in hearing research as well as objective audiometry (Somers et al., 2021). Its spatial resolution is limited by the number of electrodes that can be positioned on the scalp and the indirect measurement of neural activity through electric fields. If the goal is to postulate about the function of specific brain regions, inferring the sought activity requires knowledge about the underlying brain structures and the exact electrode locations. However, subject brain and general anatomical characteristics present a significant variability (Haładaj, 2020; Yang et al., 2019; Bartley et al., 1997). Individual head shapes also cause differences in the relative placement and studies show that the considered electrode locations can greatly affect analyses results (Hirth et al., 2020; Van Hoey et al., 2000; Wang and Gotman, 2001; Dalal et al., 2014; Schwartz et al., 1996).

While individualization of those measures is possible, every step comes at a further cost: recording the individual electrode locations after an experiment requires the appropriate hardware and additional processing time, as, depending on the method used, laborious manual processing might be necessary (Taberna et al., 2019; Koessler et al., 2007). Acquiring individual brain anatomies can come in expensive for research institutions, as neither the facilities nor the necessary resources might be available. Moreover, it is oftentimes the case that prior medical procedures or implantations prevent individuals from procedures such as magnetic resonance imaging. For hearing research specifically, cochlear implants frequently fall under the latest category (Holtmann et al., 2021; Leinung et al., 2020). In either case, a better understanding of the effects of the individualization steps may help the planning and uncertainty assessment of EEG studies.

Activity recorded by scalp-EEG sensors comprises a superposition of various brain sources, making it non-trivial to uncover underlying mechanisms. To expose specific information about the auditory functions in the brain, it is often relevant to move from the sensor-to the source space, spatially separating the signals and attributing them to their original generators. Source estimation is a complex task, involving several modelling steps. Localizing where the recorded activity actually originated from requires in the first place a representation of the elements of the subject’s head (Vorwerk et al., 2014). The scalp, skull, grey and white matter, and cerebrospinal fluid have different conductivity characteristics, requiring an appropriate model accounting for them. This information is incorporated in the forward model, which describes how the electric field generated by a cortical source is picked up as an electric potential by a sensor. Source estimation based on EEG is especially subject to errors in the forward modelling; as it is based on electric fields and the sensors are positioned directly on the skin, it is heavily influenced by the differences in conductivity estimates (von Ellenrieder et al., 2009; Leahy et al., 1998). Various solutions have been developed, and the forward-model choice mainly relies on the available computational resources and chosen measurement modality (Hallez et al., 2007; Baillet et al., 2001). For EEG research, using the boundary element method is considered an appropriate solution, offering a realistic representation of the head model (Kybic et al., 2005; Akalin-Acar and Gençer, 2004; Adde et al., 2003; Wang and Gotman, 2001). For the cases where individualization steps cannot be included, relevant software offers the option for template anatomies and electrode locations. Those can be used on the acquired experimental data, to approximate actual head characteristics.

Given the forward model, the activity of the brain regions can be estimated via the inverse solution; the sensor data is combined to create an estimate of the activity at the various brain locations. This constitutes an ill-posed problem because the number of sources is typically much larger than the number of sensors. Hence, the inverse solution is not unique and requires additional assumptions or constraints to become so (Baillet et al., 2001). Various approaches have been developed towards tackling this problem (Grech et al., 2008); among those, minimum-norm solutions fall under the category of distributed inverse solvers (Ou et al., 2009). They rely on minimal prior assumptions, and are therefore well suited in data driven approaches, where data is too noisy or no prior knowledge about source activity can be reliable (Hauk, 2004). Each grid point is considered to be the location of one or a set of equivalent current dipoles, subject to specific constraints regarding their degrees of freedom. Those algorithms look for a fitting solution to the data at each grid location simultaneously, under the restriction of a minimum overall activity amplitude. As most cognitive processing relies on distributed sources rather than isolated sources, such approaches offer an ecologically plausible solution, suited for mapping complex function in the perceptual field (Komssi et al., 2004). In the implementations of dynamic statistical parametric mapping (dSPM; Dale et al., 2000) and standardized low-resolution electromagnetic tomography (sLORETA; Pascual-Marqui, 2002), noise statistics information derived either from data or separate recordings is used to standardize the source maps, in order to compensate for depth current-orientation inhomogeneity (Hauk et al., 2011). Generally, the choice of the inverse method relies on parameters such as the sensory modality or experimental paradigm; there are, though, no precise guidelines on selecting a method, rendering the option to frequently depend on common practise and preference. Meanwhile, toolboxes offer direct implementations of multiple inverse solutions, thereby facilitating comparative studies on the same dataset, an oftentimes suggested approach (Nawel et al., 2019). In auditory research, dSPM and sLORETA are frequently applied algorithms towards solving the inverse problem (eg. Raghavan et al., 2017; Stropahl et al., 2018; Hsu et al., 2020; Mohan et al., 2020; Justen and Herbert, 2018; Jaworska et al., 2012). Based on anecdotal evidence, dSPM has been deemed to be specifically good for modeling auditory cortex sources (Stropahl et al., 2018). Instead, various method comparisons demonstrated how sLORETA can return most satisfactory results for single source localization (Grech et al., 2008).

As the first relay of auditory information, the primary auditory cortex (PAC) is essential in auditory research. Yet, with its small size and intricate placement on the superior temporal lobe (i.e. within the lateral sulcus and its non-orthogonal orientation to the scalp), the correct extraction of its activity constitutes a difficult matter (Hari and Puce, 2017). Our aim in the current study was to examine the effect of those individualization steps on the PAC localization accuracy of inverse solutions, from the experimenter’s perspective. For that reason we considered two main factors that play a crucial role in the source localization process: the electrode positions and the subject anatomy. We combined those in pairs of two, yielding four different and commonly used approaches in EEG experiments (template anatomy with template electrode positions, template anatomy with individual electrode positions, individual anatomy with template electrode positions, individual anatomy with individual electrode positions). Our basic assumption was that a fully individualized configuration should lead to the most likely precise and valid source localization (Van Hoey et al., 2000; Dalal et al., 2014; Darvas et al., 2006; Buchner et al., 1995; Akhtari et al., 1994). To control for robustness or interaction with regard to the specific inverse solutions used, we decided to conduct our study with the two inverse solutions dSPM and sLORETA.

## 2 MATERIALS AND EQUIPMENT

For the current study we analyzed data originally collected for an auditory spatial perception experiment (Baier et al., 2022). It consisted of an initial passive listening part, during which subjects were watching a silent subtitled movie. In a second part, subjects performed a spatial discrimination task on those stimuli. For our current study we only considered the EEG data during passive listening, in order to avoid any task-related characteristics.

Our dataset was recorded with a 128-channel EEG system (actiCAP with actiCHamp; Brain Products GmbH, Gilching, Germany) at a sampling rate of 1 kHz. For 23 participants we had individual anatomical structures and electrode positions. Our later event-related potential (ERP) analyses yielded missing values for three of our subjects with template attributes, hence we restricted our set to the remaining 20 subjects (9 female: mean_*age*_ = 25.4; SD_*age*_ = 2.51; 11 male: mean_*age*_ = 25.4; SD_*age*_ = 3.04).

The auditory stimuli used were complex harmonic tones (Schroeder, 1970; *F*_0_ = 100 Hz, bandwidth 1 − 16 kHz). They were presented through earphones (Etymotic Research, ER-2) and were filtered with listener-specific head-related transfer functions (HRTFs) to sound as coming from either the right or left direction on the interaural axis. The duration of every stimulus was 1.2 s with an inter-stimulus interval of 500 ms. Onset and offset ramps with raised-cosine shape had a duration of 10 ms. The stimuli were presented at a sound pressure level of about 70 dB (all three intensity offsets of 2.5, 0 and − 2.5 dB from the original study were pooled together). 600 trials were recorded per subject.

## 3 METHODS

### 3.1 EEG data preprocessing

EEG data were manually inspected to detect potential noisy channels, which were then spherically interpolated. The data were subsequently bandpass-filtered between 0.5−100 Hz (Kaiser window, *β* = 7.2, *n* = 462) and epoched ([−200, 1500] ms) relative to stimulus onset. We applied hard thresholds at − 200 and 800 *µV* to remove extremely noisy trials. Undetected bad channels were further identified through an automatic channel rejection step; if found, they would be visually inspected and interpolated. No additional noisy channels were detected for any of the subjects. We performed independent component analysis (ICA) and followed up with a manual artifact inspection and rejection of oculomotor artifacts (removal of up to 3 components per subject). The data were thereafter re-referenced to their average. Trials were equalized within each subject by pseudo-selection, in order to match the minimum amount within the subject after trial rejection and maintain an equal distribution across the recordings. On average, this resulted in 569 clean trials (SD = 27.7) per subject. All preprocessing steps were undertaken on the EEGLAB free software (Delorme and Makeig, 2004; RRID:SCR 007292).

### 3.2 Source estimation

We created four study cases, investigating the effects of different factor combinations on the source localization accuracy. The following analyses were implemented in the Brainstorm free software (Tadel et al., 2011; RRID:SCR 001761).

#### 3.2.1 Electrode locations

The first factor considered were the locations of the electrodes on a subject’s head. We examined two levels: template-based and individually tagged electrode locations. As template electrode locations we used the ICBM 152 BrainProducts Acticap 128 default EEG cap as implemented in Brainstorm, thereby matching our experimental setup. The electrode order in the channel file was first modified, to match the corresponding order of the channels in our collected data. To compare our results to those of an individualised electrode cap, we acquired the actual electrode positions through a scanning process: after data collection and while still wearing the EEG cap, an optical 3D scan (Structure Sensor with Skanect Pro, Occipital Inc., Boulder, Colorado) of each subject’s head was made, in order to capture the individual electrode position profile present during the experiment. The individual electrodes were subsequently manually tagged on the 3D scans and an electrode location file was created as an input for the upcoming head model creation steps (Fieldtrip; Oostenveld et al., 2010; RRID:SCR 004849). The default channel positions were then overwritten by the individual ones in the cases comprising individual electrode positions. Default cap locations differed from the individually tagged locations by a Euclidean distance of 0.17±0.033 mm (mean ± SD), after adjustment to the default anatomy. Similarly, after adjustment to the individual anatomy, the locations of the default cap differed from the individually tagged locations by 0.16 ± 0.035 mm.

#### 3.2.2 Subject anatomy

The second factor we examined was the subject anatomy. As in the preceding case we accounted for two levels, namely a template anatomy and an individual anatomy. The template anatomy used was the standard ICBM152 brain template as implemented in Brainstorm. For the individual subject anatomies, a structural T1-weighted magnetic resonance (MR) scan for each subject was recorded at the MR centre of the SCAN-Unit (Faculty of Psychology, University of Vienna) with a Siemens MAGNETOM Skyra 3 Tesla MR scanner (32-channel head coil; Siemens-Healthinieers, Erlangen, Germany). Anatomical MR scans for all subjects were subsequently segmented via Freesurfer (Fischl, 2012; RRID:SCR 001847), and loaded in Brainstorm. They were then used as the basis for each subject’s head model in the corresponding cases comprising individual subject anatomies. We created the anatomical models using the boundary element method in OpenMEEG (Gramfort et al., 2010; RRID:SCR 002510). Boundary surfaces were constructed by Brainstorm with 1922 vertices per layer for scalp, outer skull and inner skull, and a skull thickness of 4 mm was considered. In line with the default settings, the relative conductivity of the outer skull was set to 0.0125 and to 1 for the remaining layers. We kept the adaptive integration selected, to increase accuracy of our results. For each study and subject we performed a manual co-registration between the head models and the channel locations. In this process, the electrode cap was manually adjusted on either the template or the individual anatomy, using the translation, rotation or resizing options through the graphical user interface. To minimize individual intervention and therefore inconsistencies in reproducibility, the cap was always adjusted as a whole; no channels were fine-tuned individually. The entirely non-individualized condition with templates used for both the electrode locations and subject anatomies needed no manual co-registration, as it was already accounted for by the template models.

#### 3.2.3 Inverse solution

The third considered factor was the inverse solution used to infer cortical source activity. With dSPM and sLORETA we selected two distributed source solutions widely used and implemented in Brainstorm. Both aim for a minimum norm estimate with implicit depth weighting to improve localization accuracy of deep sources (Lin et al., 2006), but differ in the normalization approach (Nawel et al., 2019; Hauk et al., 2011). In dSPM (Dale et al., 2000), the current density normalization is done based on the noise covariance information. In sLORETA (Pascual-Marqui, 2002; RRID:SCR_013829), the current density normalization is based on the data covariance, which is a combination of the noise covariance and a modeled brain signal covariance estimate. For the calculation of the covariances, we here considered a single-trial pre-stimulus baseline interval of [− 200, 0] ms. For both solutions, the source orientations were considered constrained; in that case, a dipole, that is assumed to be placed perpendicular to the cortical surface, is considered for each vertex location (Tadel et al., 2011). Noise covariance regularization was done with a factor of *λ*^2^ = 0.1. Depth weighting and regularization parameter were selected as motivated and recommended by Brainstorm (depth weighting order = 0.5, SNR= 3 dB). Signals were reconstructed at 15000 cortical surface points.

### 3.3 Evaluation

We focused our study on the evoked activity of the right and left PAC, defined as trasversetemporal regions by the Desikan-Killiany parcellation scheme (Desikan et al., 2006). There, we evaluated the effects of the considered individualization factors with respect to both spatial and temporal aspects.

#### 3.3.1 Metrics

For each subject we extracted the evoked PAC activity for each hemisphere. These time series of current source densities were then low-pass filtered at 20 Hz (Hamming-based FIR, *n* = 150) with ERPLAB (Lopez-Calderon and Luck, 2014; RRID:SCR 009574) and baseline-corrected by a 100-ms-pre-event interval; the average across trials for each subject was subsequently calculated. Based on literature (Hari and Puce, 2017) as well as the grand average profiles, we defined a time interval for each of the signature components P1 and N1: [10 − 90] ms was defined for P1 and [50 − 150] ms for N1. In those windows, the peak amplitude (maximum for P1 and minimum for N1) and peak latency values were extracted for each component from the individual subject trial-averages, based on the `findpeaks` function as implemented in MATLAB 2018b. The single-subject data were plotted and inspected for accuracy. They were then analyzed and statistically compared based on the factors electrode location (template or individual) and subject anatomy (template or individual), individually for each hemisphere (right or left) and inverse solution (dSPM and sLORETA).

In the present study we assumed that the generating sources of P1 and N1 components are linked to focal activity in the PAC yielding maximal current source density values within it; yet localization errors likely arise, especially at neighboring vertices, due to the probability-based approach of the minimum norm estimate methods. The considered inverse solutions weigh spatially neighboring vertices higher in order to yield a smooth distribution of current source densities (Michel and Brunet, 2019). This artifact is often referred to as spatial leakage. In order to assess the leakage of localized source activity from within the PAC towards the neighboring regions, we specified a region on the cortical surface, spatially surrounding the atlas-defined PAC for each hemisphere. In order to aid reproducibility, we decided to expand the region by recruiting additional atlas-defined surrounding regions, which the PAC activity might have leaked into. As the Desikan-Killiany parcellation was deemed too coarse, we based our new region selection on the finer Destrieux atlas (Destrieux et al., 2010). We hence constructed an extended region of interest (ROI) by merging the regions of the planum temporale, fissure, transverse temporal sulcus, circular sulcus as well as our initially defined PAC region. On the right hemisphere, the area covered by the extended ROI spanned 24.67 cm^2^ versus the 4.56 cm^2^ of the initially defined PAC region (factor of 5.4). On the left side, the original PAC surface of 6.16 cm^2^ was expanded to 28.03 cm^2^ (factor of 4.6). For each of the two components (P1 and N1) we considered the previously extracted peak latency found for each subject average; for the exact same time points we extracted the activity of the extended ROI. We then calculated the squared amplitude ratio between the two [(summed activity over PAC vertices)^2^/(summed activity over extended ROI)^2^], denoted as ”ROI power ratio”. This metric quantifies the proportion of evoked power contained in the PAC relative to that occurring in the extended ROI and is thus assumed to reflect the leakage to the neighboring regions in the sense that higher ratios indicate less leakage.

All aforementioned analysis procedures were implemented in MATLAB 2018b (RRID:SCR 001622).

#### 3.3.2 Statistical analysis

Statistical analyses on the source localized time series and the extracted data relied on a mixed-model design with a multi-way ANOVA, considering a within-subject design. In particular, the analysis of the peak amplitude, peak latency, and ROI power ratio included two factors with two levels each: subject anatomy (template or individual) and electrode positions (template or individual). All ANOVAs were performed separately for each inverse solution and hemisphere.

Before each test, data were z-scored within subject and transformed according to the Box-Cox transformation (Hawkins and Weisberg, 2017). Furthermore, we ran Levene’s test assessing violations in the homogeneity of variance and inspected the ANOVA residuals verifying the normality assumption. Post-hoc analyses of interactions/contrast have been done with Bonferroni correction. Finally, for the metrics that violated these assumptions, non-parametric aligned ranks transformation ANOVA (Wobbrock et al., 2011) was performed and the Wilcoxon test was used in the post-hoc analysis. Effect sizes are only reported if the parametric ANOVA was applied.

All statistical analyses were performed in R Project for Statistical Computing (RRID:SCR 001905). In addition to the standard environment, we relied on the following packages for the analysis: `afex` for the ANOVA tests (Barr et al., 2013), `emmeans` for the posthoc comparison (RRID:SCR 018734), `ARTool` for the non-parametric ANOVA (Wobbrock et al., 2011) and `ggplot2` for data visualization (RRID:SCR 014601).

## 4 RESULTS

### 4.1 Evoked PAC activity

We compared average event-related PAC responses to sounds locked to the stimulus onset. Figure 1 shows time courses comparing the different individualization levels for the two hemispheres and inverse solutions. For all source localization conditions we reconstructed stereotypical auditory-evoked responses in the PAC with a prominent positive deflection between 10 and 90 ms, denoted as the P1 component, followed by a negative deflection between 50 and 150 ms, denoting the N1 component. The left hemisphere (figure 1A, C) was more clearly affected by the different configurations than the right hemisphere (figure 1B, D). Later components were moreover more susceptible to latency differences than earlier components. Additionally, time series profiles differed depending on the inverse solution used; dSPM curves (figure 1A, B) appeared more pronounced for the fully individualized configurations, while template peaks were rather more salient among the sLORETA curves (figure 1C, D). We statistically analyzed the extracted source activation profiles for each hemisphere and inverse solution separately. Detailed information regarding the extracted values can be found in the supplementary material.

**Figure 1.**
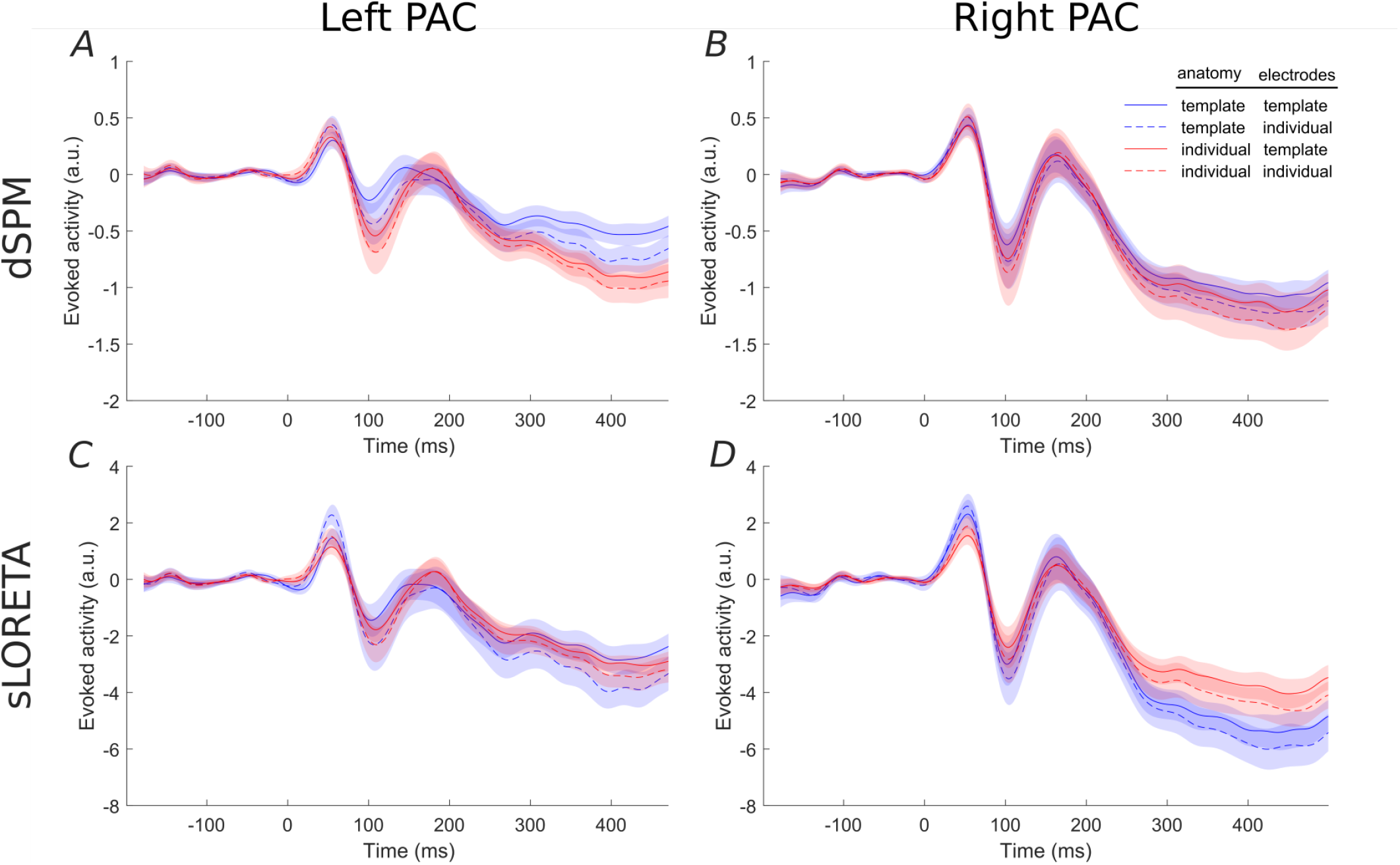
Evoked PAC activity inferred via dSPM (top) and sLORETA (bottom) for the left (A, C) and right (B, D) PAC. Shaded areas denote the s.e.m. values. Visualization of the averaged attributed values (i.e. peak’s amplitude and latency) over the different conditions is available in the Supplementary Material.

Based on dSPM, the left PAC (figure 1A) shows a differentiation depending on the degree of individualization. At P1, peak latencies were shorter for individual-than template electrode locations (*F* = 6.59, *p* = 0.01) and peak amplitudes increased with individual electrodes locations only within template anatomies (*F* = 12.16, *p* < 0.001). At later time points, the characteristics appear to be driven by the inclusion or not of a template or individual brain anatomy. Concordantly, only the use of individual anatomy yielded a significant increase of the N1 amplitude (*F*_1,19_ = 5.31, *p* = 0.03, *η*^2^ = 0.22). N1 latencies were significantly longer for individual anatomies (*F*_1,19_ = 16.17, *p* < 0.001, *η*^2^ = 0.46) and individual electrode locations (*F*_1,19_ = 17.19, *p* < 0.001, *η*^2^ = 0.47).

In the right PAC (figure 1B) the curves of all four individualization conditions are highly overlapping, suggesting no strong impact of any of the individualization steps. Nevertheless, individual electrode locations resulted consistently in slightly larger P1 amplitudes (*F* = 16.69, *p* < 0.001).

For the N1 component, electrode locations were again a significant factor (*F* = 6.99, *p* < 0.01), yielding more pronounced peaks with individualization. No significant effect was found on either the P1 or N1 latencies.

sLORETA applied to the left hemisphere (figure 1C) revealed a significant interaction between anatomy and electrode locations on the P1 amplitude (*F* = 11.62, *p* < 0.001); individual electrode locations generated highest values in particular when combined with template anatomies. Significant differences were neither found on P1 latencies nor on N1 amplitudes. The N1 latency, though, was affected by both anatomy (*F* = 15.45, *p* < 0.001, *η*^2^ = 0.46) and electrode locations (*F* = 11.86, *p* < 0.01, *η*^2^ = 0.40); overall individual anatomies produced later N1 peaks than the corresponding template configurations and individual electrode locations yielded later peaks.

On the right hemisphere (figure 1D), anatomy (*F*_1,19_ = 10.31, *p* < 0.001, *η*^2^ = 0.35) and electrode locations (*F*_1,19_ = 20.38, *p* < 0.001, *η*^2^ = 0.52) showed significant main effects on P1 amplitude for sLORETA. Template anatomies produced higher peak values, more so in combination with individual electrode locations. The interaction between anatomy and electrode locations was found significant for P1 latency (*F*_1,19_ = 5.22, *p* < 0.05, *η*^2^ = 0.22); the combination of individual electrode locations and template anatomies yielded the shortest values. Anatomy caused a differentiation on the N1 peak values (*F* = 7.05, *p* < 0.01), where template anatomies led to slightly more pronounced peaks. No significant effects were found on N1 latency.

### 4.2 Spatial leakage

We inferred the brain activity not only to the PAC but to the entire cortical surface. The corresponding brainmaps are shown in figure 2 for dSPM and figure 3 for sLORETA. There was a clear evoked activation in the temporal region, yet that activation differed in its precise location and spread, depending on the particular configuration considered. Generally, the activation was attributed to regions extending rather more superior and posterior than the atlas-defined PAC (cyan outline) and the overall pattern seemed to be dominated by the subject anatomies. The configurations comprising individual subject anatomies exhibited more constrained activation patterns within the atlas-defined PAC, whereas those with the template anatomies showed a higher spread of activation. This was most pronounced within the extended ROI area surrounding and including the PAC but also reached further to other parts of the temporal and parietal lobes. When regarding the general brain profiles, electrode locations appeared to play a secondary role; individual electrode locations did not consistently seem to constrain the activation further. Between the two inverse solutions, the spread was noticeably more distributed in the use of sLORETA, compared to dSPM.

**Figure 2.**
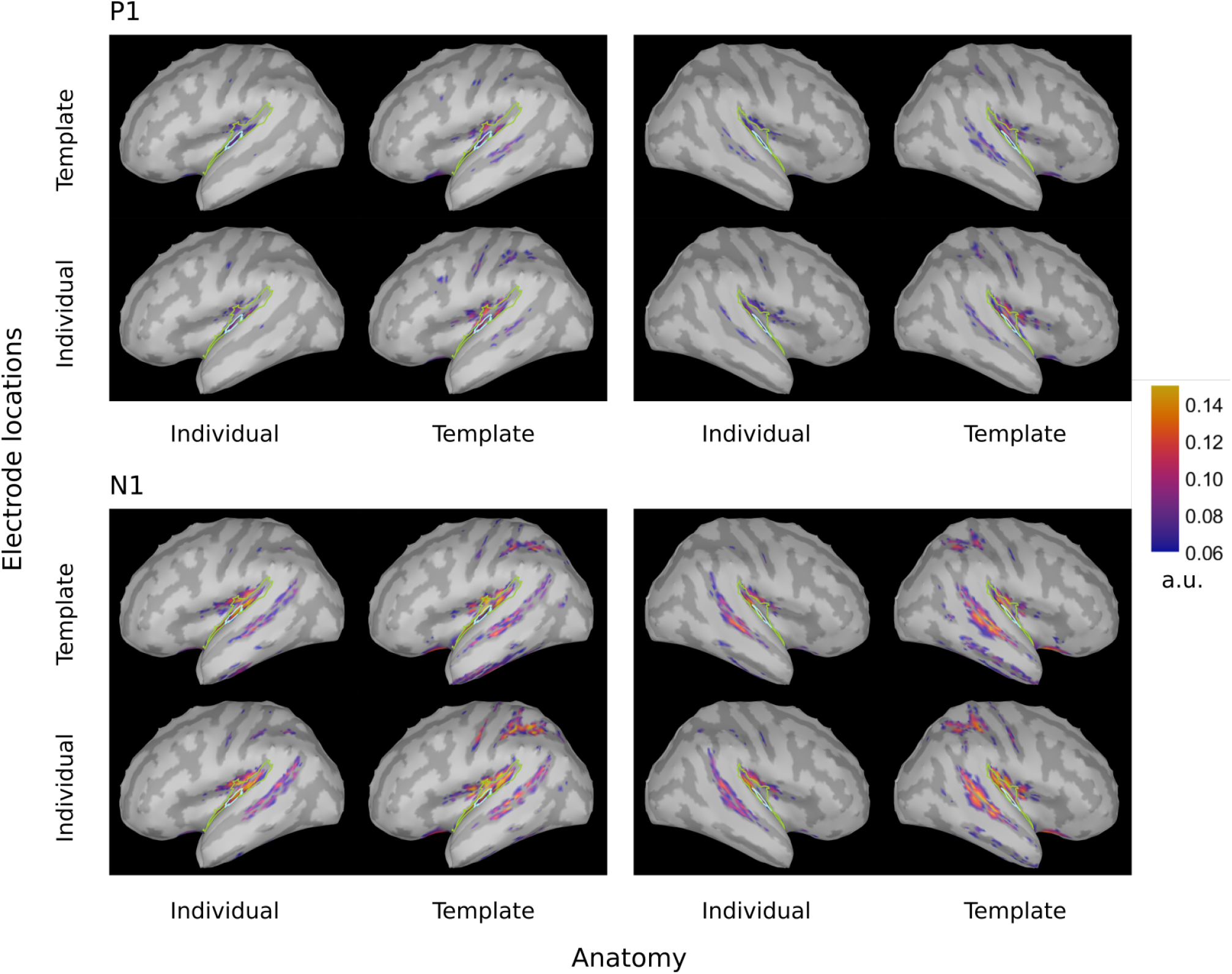
dSPM brainmaps depicting the PAC activation at the P1 (∼ 55 ms, top) and N1 (∼ 100 ms, bottom) timing for both hemispheres and all considered configurations. Cyan: atlas-defined PAC, green: extended ROI

**Figure 3.**
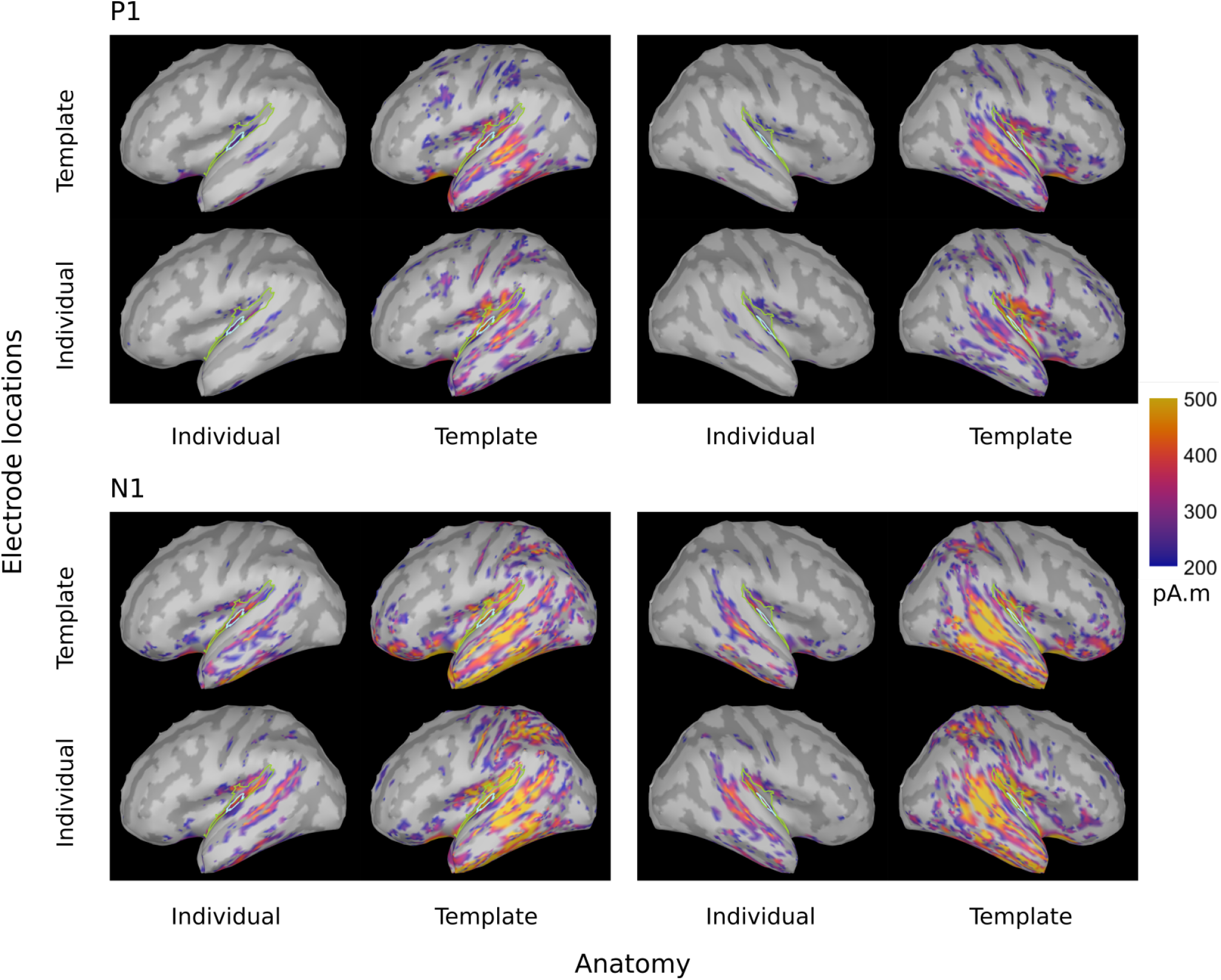
sLORETA brainmaps depicting the PAC activation at the P1 (∼ 55 ms, top) and N1 (∼ 100 ms, bottom) timing for both hemispheres and all considered configurations. Cyan: atlas-defined PAC, green: extended ROI

Complementary to the brainmaps, we evaluated the P1 and N1 power ratios between the PAC and the extended ROI (figure 4). When considering the dSPM inverse solution and the left hemisphere, individualization of the electrode locations led to higher P1 ratio (*F*_1,19_ = 5.27, *p* < 0.05, *η*^2^ = 0.22). No significant effects were found on the corresponding ratio for the N1 component.

**Figure 4.**
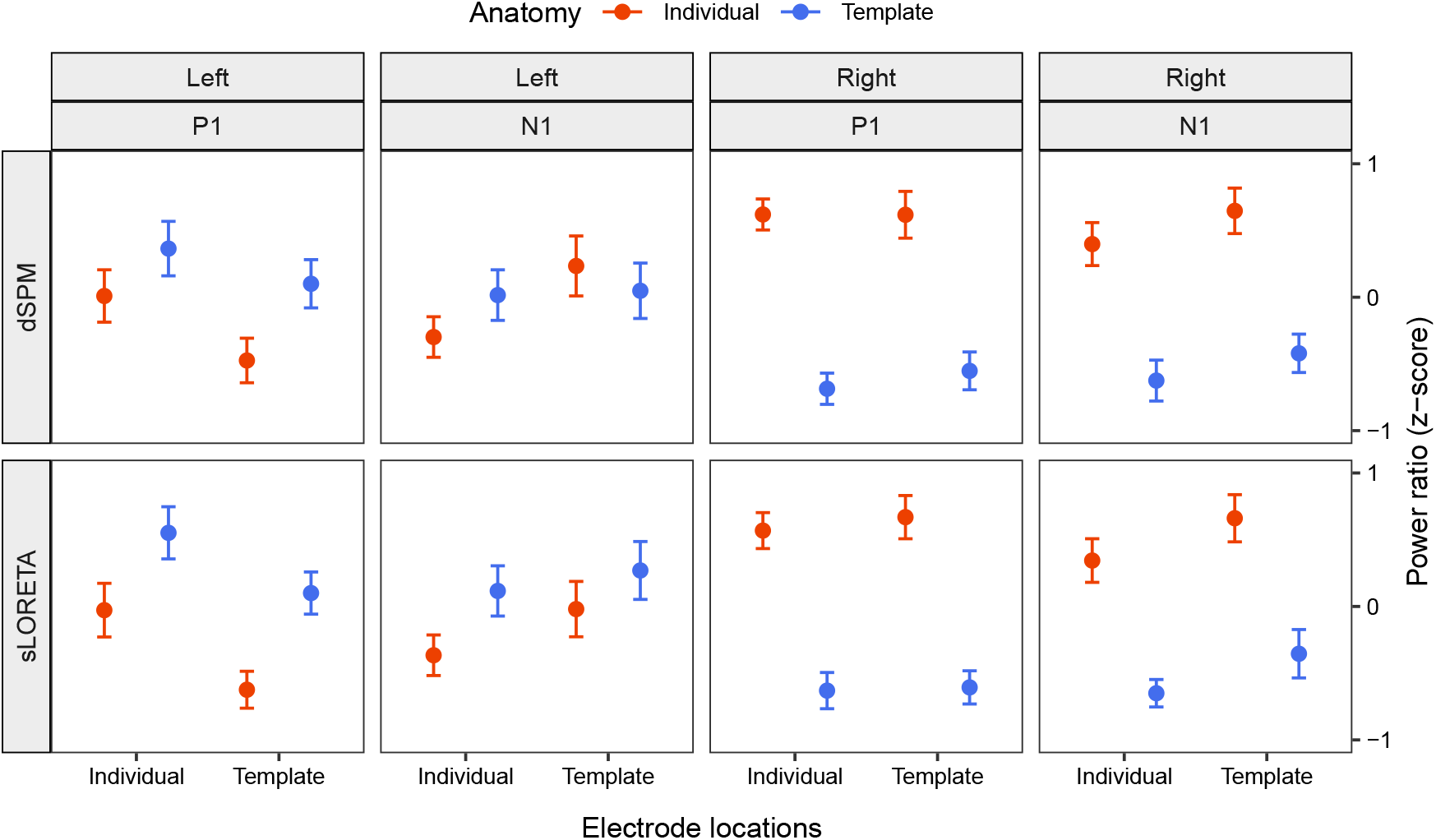
ROI power ratios for P1 and N1 components inferred via dSPM (top) and sLORETA (bottom) for the right and left hemisphere. Higher ratios are interpreted as indication for lower spatial leakage of inferred PAC source activity. Values correspond to the average over subjects after z-scoring per hemisphere, inverse solution and subject. Error bars denote the s.e.m. values.

In the right hemisphere under the dSPM inverse solution, anatomy had a significant main effect on both the P1 (*F* = 71.14, *p* < 0.001) and N1 (*F*_1,19_ = 21.84, *p* < 0.001, *η*^2^ = 0.53) power ratios. Individual anatomies generated higher power ratios at the peak of both components.

When considering the sLORETA inverse solution in the left hemisphere, anatomy (*F*_1,19_ = 7.05, *p* < 0.05, *η*^2^ = 0.27) and electrode locations (*F*_1,19_ = 9.51, *p* < 0.01, *η*^2^ = 0.33) were main effects for the P1 power ratio. Individual subject anatomies led to lower power ratio values, while individual electrode locations ameliorated the result. The corresponding N1 power ratios were significantly affected only by anatomy (*F* = 4.94, *p* < 0.05), with individualized subject anatomies leading to a deterioration of the value, hence denoting higher leakage.

In the corresponding right hemisphere, anatomy was a significant main effect for both P1 (*F* = 69.92, *p* < 0.001) and N1 (*F*_1,19_ = 17.64, *p* < 0.001, *η*^2^ = 0.48). In both cases individualization benefited the localization accuracy.

## 5 DISCUSSION

In the present study our aim was to single out the effects of different individualization steps on the accuracy of inferring PAC activity from EEG data. We compared combinations of template or individualized electrode locations and subject anatomies while using two different inverse solutions (dSPM and sLORETA). Through that we reconstructed and characterized the evoked PAC time series and assessed the spatial leakage around the PAC in each hemisphere. Table 1 summarizes the significant effects for all configurations and their consistency with individualization benefit. As evident, both the factors of electrode location as well as subject anatomy were found to have an impact on the defined current source density metrics; yet their effect varied depending on the target brain area (PAC in the left or right hemisphere), the evoked component characteristic (amplitude or latency of P1 or N1), and also the type of inverse solution (dSPM or sLORETA) considered.

**Table 1.**
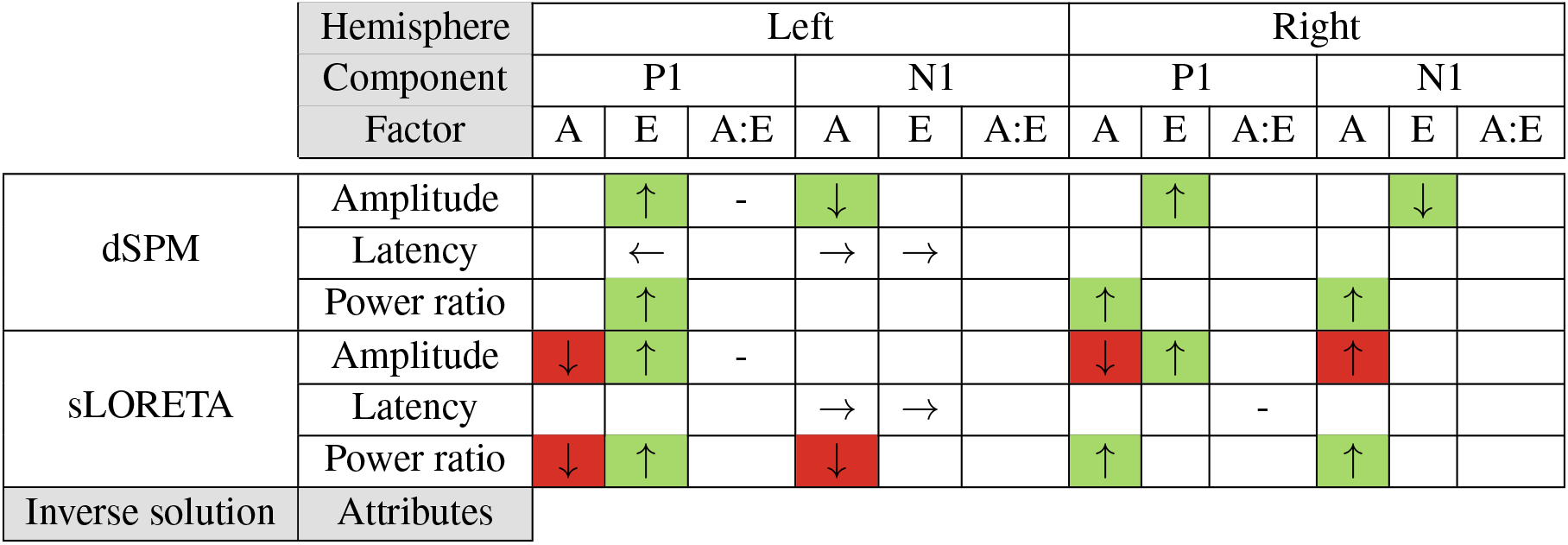
Summary of the effects of anatomy (A) and electrode locations (E) as well as their interaction (A:E) resulting from the statistical tests. Only significant results are reported (*p* < 0.05). Arrows indicate the direction of change from the template factor level to the individual factor level. Dashes indicate opposing interactions. Color coding denotes the assessment of every such change as individualization benefit (green) or degradation (red). If this interpretation on the direction of change seemed ambiguous, as is the case for latency changes and opposing interactions, the cell has not been colored.

|  | Hemisphere<br>Component<br>Factor | Left |  |  |  |  |  | Right |  |  |  |  |  |
| --- | --- | --- | --- | --- | --- | --- | --- | --- | --- | --- | --- | --- | --- |
|  |  | P1 |  |  | N1 |  |  | P1 |  |  | N1 |  |  |
|  |  | A | E | A:E | A | E | A:E | A | E | A:E | A | E | A:E |
| dSPM | Amplitude |  | ↑ | - | ↓ |  |  |  | ↑ |  |  | ↓ |  |
|  | Latency |  | ← |  | → | → |  |  |  |  |  |  |  |
|  | Power ratio |  | ↑ |  |  |  |  | ↑ |  |  | ↑ |  |  |
| sLORETA | Amplitude | ↓ | ↑ | - |  |  |  | ↓ | ↑ |  | ↑ |  |  |
|  | Latency |  |  |  | → | → |  |  |  | - |  |  |  |
|  | Power ratio | ↓ | ↑ |  | ↓ |  |  | ↑ |  |  | ↑ |  |  |
| Inverse solution | Attributes |  |  |  |  |  |  |  |  |  |  |  |  |

### 5.1 Individualization factors

The considered individualization factors influenced the two hemispheres quite differently. In the right hemisphere, anatomy affected mainly the power ratio indicating spatial leakage, while electrode positions had an impact on peak amplitudes. Contrary to that, we found a more complex pattern of effects in the left hemisphere: individualized solutions gave earlier peaks for P1 and later ones for N1, individual electrode locations increased both the P1 amplitude and power ratio, and individual anatomies interacted with that effect on P1 amplitude and independently enlarged N1 amplitudes.

We speculate that such a regional variance in source reconstruction could be resulting from either state or trait effects. On the one hand, source localization estimates could show higher variance across subjects because of brain morphology (i.e., cerebral size, Bartley et al., 1997; handedness, Good et al., 2001) that, on a group level, may result in higher or lower uncertainty for different regions, especially for template solutions. Higher inter-individual variability is also generally found in the left auditory cortex (Ren et al., 2021). On the other hand, the observed asymmetry might be related to stimulus features; auditory stimulation characteristics preferentially processed on either hemisphere, such as unattended automatic change detection of spectral or temporal features, may influence the variance of the source reconstruction estimates (Okamoto et al., 2009; Schönwiesner et al., 2007). Our stimuli moreover carry spatial characteristics, as they are presented from either the left or right side of- and at different distances from the listener. Spatial processing has been shown to exhibit right-hemispheric dominance, possibly explaining the differences we observe (Deng et al., 2020; Middlebrooks, 2015; Kaiser et al., 2000).

Individual anatomies and electrode locations allow for a more precise attribution of the recorded activity to the corresponding regions, thereby likely accounting for the various inter-individual variability characteristics. Acquiring individual electrode locations, though, usually comes with considerable measurement uncertainty; to some degree this also depends on the acquisition strategy. With our procedure, a considerable amount of the experimenter’s individual intervention is necessary in obtaining the 3D scan as well as tagging the electrodes. The extent of it might differ, when more automatized - and therefore also more reproducible - methods are used (Hirth et al., 2020; Koessler et al., 2011), potentially yielding different effects regarding the choice between template or individual electrode locations.

As we were interested in localizing the PAC we restricted our search on the cortical surface and this is where our results apply. Our choice of constrained sources in the brain might be an essential contributor to our outcomes: when individual anatomies are unavailable, selecting fixed sources might be too restricting and introduce errors in the considered orientations (Westner et al., 2022; Hillebrand and Barnes, 2003); therefore a different setting might be more suited for the case of template anatomies.

In order to extract the targeted cortical activity, we focused on the PAC region as defined by the Desikan-Killiany atlas. Yet, as seen on figure 2 and 3, none of the configurations seem to perfectly capture the core of the PAC activation. Different atlases vary in their parcellation; as a result, using a different parcellation scheme for such an investigation might capture the activation differently and hence lead to deviating results regarding the accuracy of localization. Another possibility could be to move away from an atlas-based- and towards a functional ROI definition. Extending the study by manually defining the PAC based on the observed, and therefore actual source activation, could increase the effect sizes and give further insight into the localization precision offered by the individualization steps.

Given the aforementioned limitations and despite the clear loss in spatial acuity obtained without individualization, we found the stereotypical auditory evoked response elicited within the PAC in all configurations (figure 1). In that regard the EEG based source localization of early evoked activity may be considered satisfactory in all cases. This is important especially in occasions where no individualization steps can be taken, as could happen with infants, implantees or situations where the corresponding resources (time, personnel or funds) are not available. Contrarily, there are cases where individualization is indispensable. Such can be investigations with known underlying structural differences, as could be in the case of hearing loss (Chen et al., 2021; Manno et al., 2021; Alfandari et al., 2018). In a general setting, though, where no such restrictions apply, our results can aid in the direction of designing the aspects of an experimental study: depending on the effect examined and available resources, decisions can be made about whether template configurations would be sufficient or a further individualization, whether electrode locations or subject anatomy, would be in order.

### 5.2 Differences between inverse solutions

Regarding the choice of an inverse solution itself, different algorithms are based on different prior assumptions (Grech et al., 2008). We here restricted our study to two widely used methods falling under the same algorithmic category (minimum-norm solutions); an informative and oftentimes suggested way is to compare different algorithms before drawing conclusions on the plausibility of the results (Nawel et al., 2019).

When comparing the analyses outcomes of our considered source localization configurations as shown in table 1, the differences between inverse solutions become noticeable. With sLORETA individualization steps yielded rather inconsistent main effects: we found some benefit of individual electrode positions, yet anatomy seemed to work in the opposite direction than what was expected. Inclusion of individual subject anatomies had incongruent effects on our metrics, oftentimes leading to a deterioration of the accuracy with higher individualization (table 1, red cells). Contrary to that, dSPM showed consistent results: all main effects contributed towards an amelioration of the considered values with individualization of either the subject anatomy or the electrode positions.

Though not reflected in the metrics of table 1, there is a considerable difference in the overall activity spread between dSPM and sLORETA. The activity is largely distributed over the temporal and parietal lobes using sLORETA, while with dSPM it remains rather spatially focused. As such, dSPM seems to be more suitable for capturing more focal auditory processes targeting specific regions implicated in them.

In sum, our findings demonstrate the benefit of using additional individualized information regarding brain anatomy and electrode positioning; they further support previous notions towards using dSPM for investigating auditory processes (Stropahl et al., 2018). A restricted activation profile can be especially beneficial, for instance, when considering a differentiation between the ventral and dorsal auditory stream, both of which also comprise relatively small areas (Bizley and Cohen, 2013).

## AUTHOR CONTRIBUTIONS

KI and RBau designed the study. KI performed the EEG analyses. RBar performed the statistical analyses. KI, RBau, and BT interpreted the results. KI and RBar wrote the first draft of the manuscript and all authors revised it.

## SUPPLEMENTAL DATA

Amplitude values for P1 and N1 components as well as their latencies are provided in the supplementary material, submitted with the present manuscript.

## DATA AVAILABILITY STATEMENT

Data are available by the authors upon reasonable request.

