## Supplementary Figures for "Benefits of individualized brain anatomies and EEG electrode positions for auditory cortex localization"

### *Supplementary Material*

#### 1 SUPPLEMENTARY TABLES AND FIGURES

##### 1.1 Figures

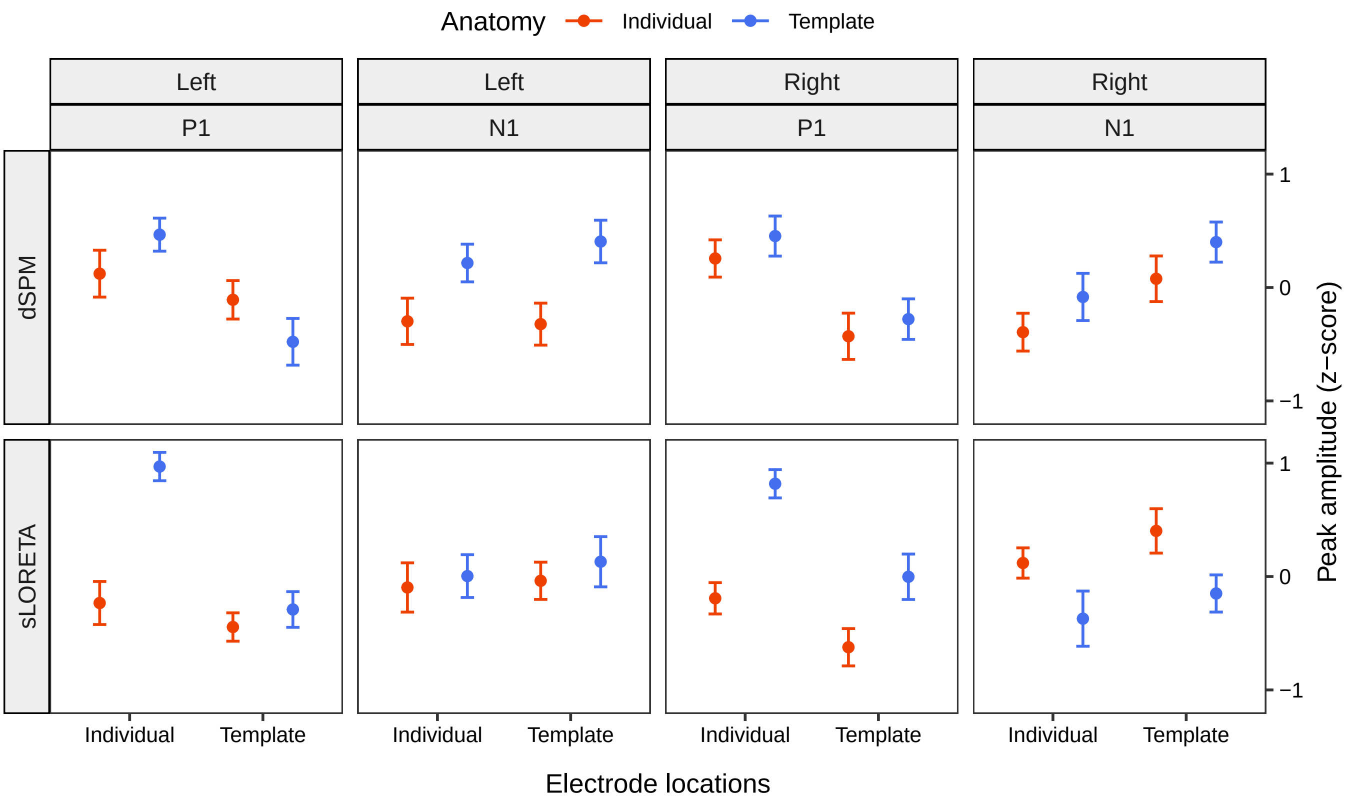


**Figure S1.** Peak PAC amplitudes for P1 and N1 components inferred via dSPM (top) and sLORETA (bottom) for the right and left hemisphere. Values correspond to the average over subjects after z-scoring per condition and within subject. Error bars denote the s.e.m. values.

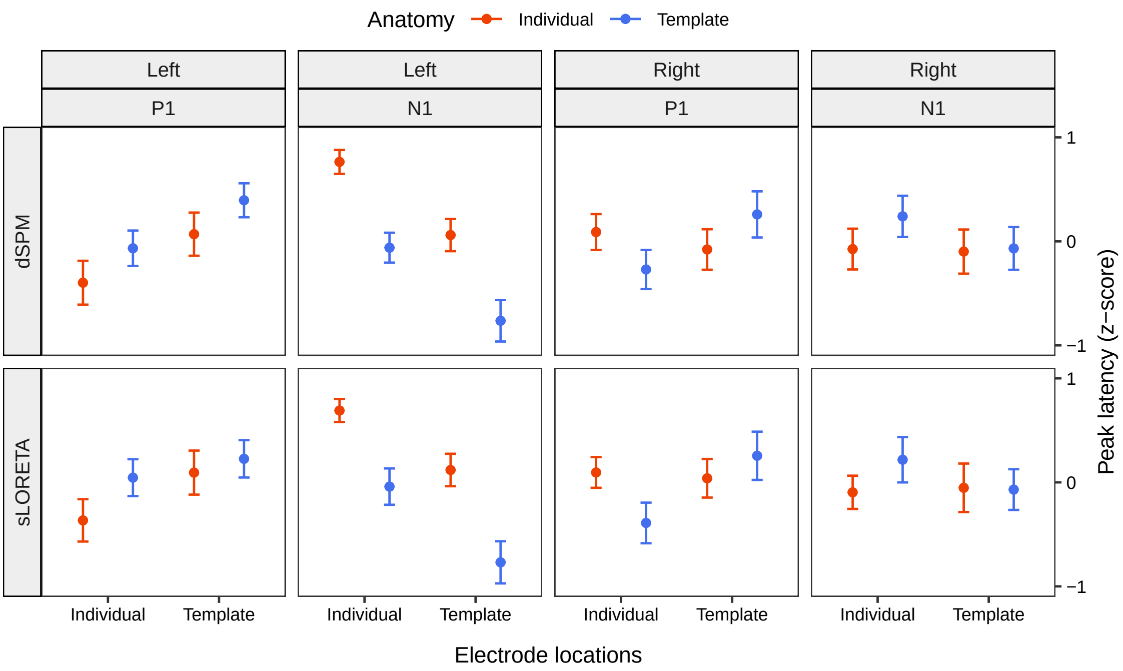


**Figure S2.** Latencies for P1 and N1 components inferred via dSPM (top) and sLORETA (bottom) for the right and left hemisphere. Values correspond to the average over subjects after z-scoring per condition and within subject. Error bars denote the s.e.m. values.
